# An ultrapotent neutralizing bispecific antibody with broad spectrum against SARS-CoV-2 variants

**DOI:** 10.1101/2021.08.10.455627

**Authors:** Hui Zhang, Haohui Huang, Rong Li, Lu Zhang, Zhiwei Wang, Jiaping Li, Junyou Chen, Huafei Su, Dandan Zheng, Ziqi Su, Li Wang, Chunping Deng, shujun Pei, Shenghua Zhu, Chan Li, Yaochang Yuan, Haitao Yue, Yanqun Wang, Xiaobo Li, Cuihua Liu, Jinchen Yu, Hui Zhang, Shengfeng Li, Xianming Huang

## Abstract

In spite of the successful development of effective countermeasures against Covid-19, variants have and will continue to emerge that could compromise the efficacy of currently approved neutralizing antibodies and vaccines. Consequently, novel and more efficacious agents are urgently needed. We have developed a bispecific antibody, 2022, consisting of two antibodies, 2F8 and VHH18. 2F8 was isolated from our proprietary fully synthetic human IDEAL (Intelligently Designed and Engineered Antibody Library)-VH/VL library and VHH18 is a single domain antibody isolated from IDEAL-nanobody library. 2022 was constructed by attaching VHH18 to the C-terminal of Fc of 2F8. 2022 binds two non-overlapping epitopes simultaneously on the RBD of the SARS-CoV-2 spike protein and blocks the binding of RBD to human angiotensin-converting enzyme 2 (ACE2). 2022 potently neutralizes SARS-CoV-2 and all of the variants tested in both pseudovirus and live virus assays, including variants carrying mutations known to resist neutralizing antibodies approved under EUA and that reduce the protection efficiency of current effective vaccines. The half-maximum inhibitory concentration (IC50) of 2022 is 270 pM, 30 pM, 20 pM, and 1 pM, for wild-type, alpha, beta, and delta pseudovirus, respectively. In the live virus assay, 2022 has an IC50 of 26.4 pM, 13.3 pM, and 88.6 pM, for wild-type, beta, and delta live virus, respectively. In a mouse model of SARS-CoV-2, 2022 showed strong prophylactic and therapeutic effects. A single administration of 2022 intranasal (i.n.) or intraperitoneal (i.p.) 24 hours before virus challenge completely protected all mice from bodyweight loss, as compared with up to 20% loss of bodyweight in placebo treated mice. In addition, the lung viral titers were undetectable (FRNT assay) in all mice treated with 2022 either prophylactically or therapeutically, as compared with around 1×10^5^ pfu/g lung tissue in placebo treated mice. In summary, bispecific antibody 2022 showed potent binding and neutralizing activity across a variety of SARS-CoV-2 variants and could be an attractive weapon to combat the ongoing waves of the COVID-19 pandemic propagated mainly by variants, especially, the much more contagious delta variant.

## Introduction

The COVID-19 pandemic is ongoing continuously around the world. As of August 2021, there are over 200 million confirmed cases and 4 million deaths across nearly 200 countries (https://COVID19.who.int/). The SARS-CoV-2 virus has continued to evolve throughout the course of the pandemic (1-3). Variants continue to emerge, and some variants might become more contagious, more virulent, or more resistant to the current effective vaccines and neutralizing antibodies, which are often referred to as variants of concern (VOC) (2, 4, 5). There currently are 4 VOCs named by WHO, including alpha, beta, gamma, and delta.

Several neutralizing antibodies have demonstrated clinical efficacy, and were approved by the FDA under an Emergency Use Authorization (EUA) for the treatment of patients with mild to modest COVID-19, especially for those with pre-existing medical conditions, who are at higher risk of developing severe symptoms following infection (6-8). Numerous neutralizing antibodies are currently in advanced clinical development and have shown promising efficacy. As the COVID-19 pandemic continues to spread, neutralizing antibodies, as a readily available therapeutic option, play an important role in the fight against COVID-19, effectively protecting vulnerable people from getting infected or from developing severe diseases following SARS-CoV-2 infection (9, 10).

Most of the neutralizing antibodies target the RBD, the critical functional domain responsible for viral infection of host cells (11, 12). Unfortunately, mutations are inevitable for the SARS-CoV-2 coronavirus, just like many other viruses. Variants have emerged carrying mutations in the RBD that could potentially weaken the effectiveness of existing neutralizing antibodies or vaccines (13-15). Studies have consistently shown that some mutations in the RBD, such as K417N/T, N439I, L452R, E484K/Q, and N501Y, can substantially reduce the strength of some neutralization antibodies and vaccines (15-17). For instance, FDA has revoked the EUA for Bamlanivimab monotherapy for COVID-19 due to the loss of efficacy against some of the currently circulating VOCs. Diamond and colleagues have recently shown that REGN10933, LY-CoV555, and 2B04 exhibited a marked or complete loss of neutralizing activity against variants B.1.351, B.1.1.28, and viruses containing the E484K mutation (18). Furthermore, studies have shown that monotherapy with a single antibody could lead to virus escape, both in vitro and in vivo (19-21). Indeed, the current resurgence at an alarming speed of the COVID-19 pandemic in regions with near herd immunity has generated major concern around the world with numerous cases of breakthrough infections (22-24). Therefore, novel antibodies that maintain their neutralization strength and breadth against the abovementioned resistant variants and possibly future emerging variants are urgently needed, which could offer extremely valuable and readily available countermeasures to combat the current wave of the COVID-19 pandemic caused mainly by VOCs.

In this study, we developed 2022, a fully-human bispecific antibody, that binds two distinct epitopes on the RBD simultaneously, blocks the interaction between the RBD and ACE2, and potently neutralizes all currently (as of August 4, 2021) known VOCs of SARS-CoV-2, such as alpha, beta, gamma, and delta, including variants carrying mutations known to be resistant to current effective countermeasures. 2022 could be a very attractive and potent therapeutic to combat the current COVID-19 pandemic caused by the troublesome VOCs.

## Results

### Characterization of monoclonal neutralizing antibody 2F8 and VHH18

Spike RBD is the major target for SARS-CoV-2 neutralizing antibodies. To isolate neutralizing antibodies against SARS-CoV-2, we used recombinant RBD of SARS-CoV-2 spike protein as antigen, panning from Bio-Thera’s proprietary IDEAL synthetic VH/VL and nanobody fully human phage library separately. After four rounds of panning, soluble scFv or nanobody candidates were first assessed for RBD binding in ELISA, positive clones were then converted to full IgG or VHH-Fc and assessed for their abilities to block RBD binding to ACE2 and to neutralize SARS-CoV-2 pseudovirus, and for epitope binning. The two best lead antibodies 2F8 and VHH18, isolated from the VH/VL library and nanobody library, respectively, were chosen to generate the bispecific antibody, 2022. Both 2F8 and VHH18 bind specifically to recombinant RBD and the ectodomain of trimeric spike of SARS-CoV-2 with high affinity in ELISA, with EC50 values of 130 pM and 150 pM for RBD (Figure 1a), 130 pM and 190 pM for spike trimer (Figure 1b), respectively; in addition, 2F8 can completely block the binding of RBD to recombinant human ACE2, with an IC50 value of 190 pM; interestingly, VHH18 can only partially block the interaction of RBD with recombinant human ACE2 (Figure 1c). Furthermore, both 2F8 and VHH18 can potently neutralizing SARS-CoV-2 pseudovirus, with IC50 values of 45 pM and 1600 pM, respectively (Figure 1d). Importantly, 2F8 and VHH18 bind distinct non-overlapping epitopes on RBD as determined by Fortebio in-tandem assay. When immobilized RBD was first loaded with saturated 2F8, VHH18 was still able to bind RBD completely, indicating that 2F8 and VHH18 can bind RBD simultaneously and 2F8 and VHH18 bind distinct epitopes on RBD (Figure 1e).

**Figure. 1:**
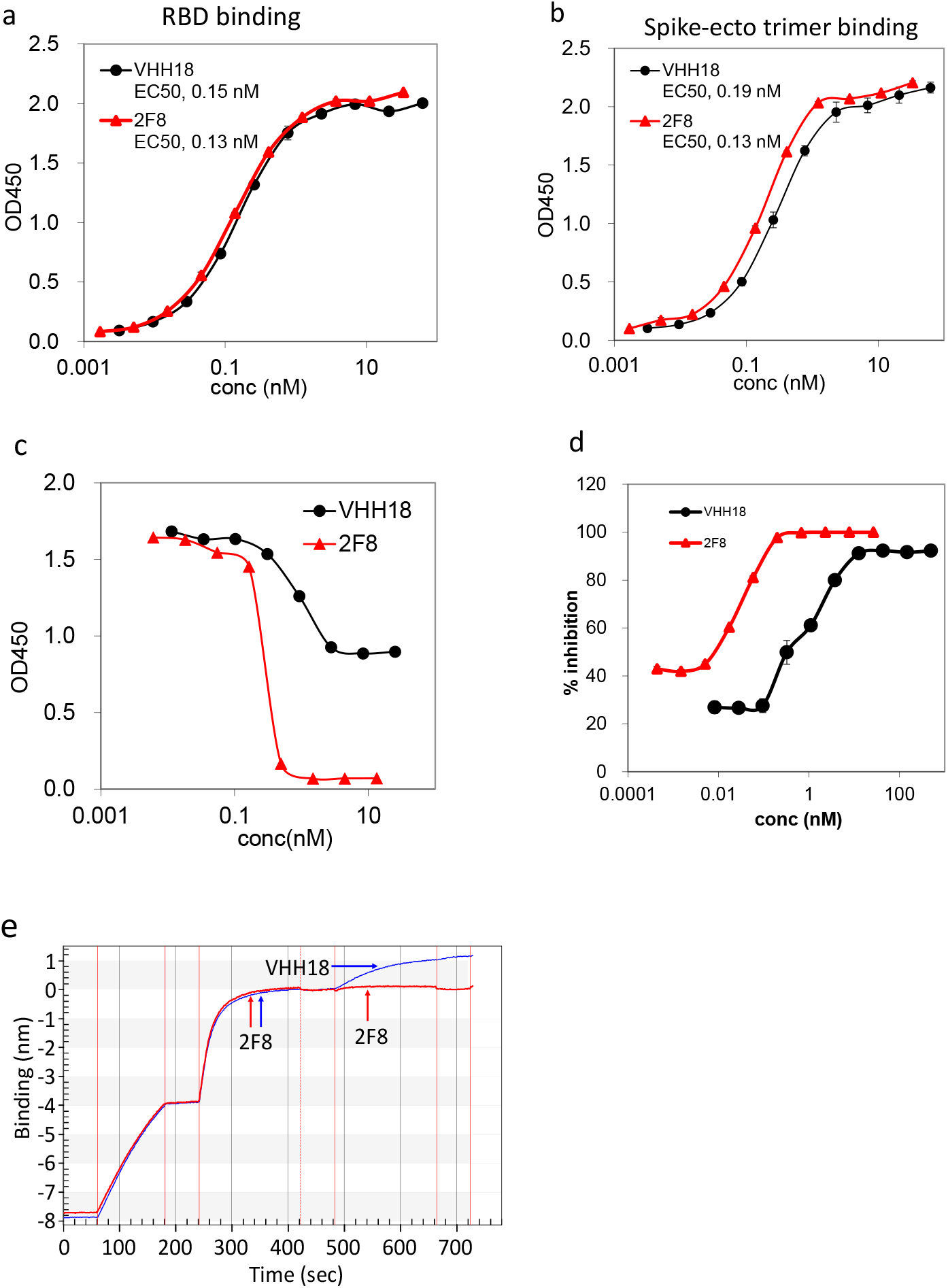
Characterization of 2F8 and VHH18 in vitro. a. Binding of 2F8 and VHH18 to recombinant RBD in ELISA. Data are mean ± S.D. of technical duplicates. EC50 values are shown on the graph. b. Binding of 2F8 and VHH18 to recombinant spike ectodomain trimer in ELISA. Data are mean ± S.D. of technical duplicates. EC50 values are shown on the graph. c. Blocking of RBD binding to recombinant human-ACE2 in ELISA. d. Neutralization activity against SARS-CoV-2 pseudovirus. e. Epitope binning of 2F8 and VHH18 by in-tandem ForteBio. RBD was immobilized on chip followed by saturated 2F8, subsequently, VHH18 or 2F8 was loaded. VHH18 binds RBD in the presence of saturated 2F8, indicating that VHH18 and 2F8 have distinct epitopes on RBD.

### Binding characterization of bispecific neutralizing antibody 2022

Cocktails of neutralizing antibodies against SARS-CoV-2 have been successfully applied to combat viral escape in clinical (6, 25). A bispecific antibody can combine the advantages of cocktails into a single molecule. We chose 2F8 and VHH18 to generate the bispecific neutralizing antibody 2022. As described above, 2F8 and VHH18 have non-overlapping distinct epitopes on RBD and both show strong neutralizing activity against SARS-CoV-2. 2022 was generated by linking VHH18 to the C-terminal of the Fc of 2F8 (Figure 2a). The binding activity of 2022 was first assessed by ELISA. As shown in Figure 2b, 2022 bound strongly to both RBD and spike-trimer of SARS-CoV-2, with EC50 value of 270 pM and 260 pM, respectively. Most neutralizing antibodies inhibit SARS-CoV-2 infection by blocking the binding of RBD to ACE2. The activity of 2022 to inhibit the binding of RBD to ACE2 was assessed by ELISA. As shown in Figure 2c, 2022 strongly inhibited binding of ACE2 to RBD (IC50=180 pM). 2022 was then evaluated for binding to occurring SARS-CoV-2 variants tested, including mutations such as K417N/T, E484K, N501Y, L452R, and D614G, carried by variants of concern such as alpha, beta, gamma, and delta, with IC50 values ranging from 200 to 280 pM, indicating that these mutations in the RBD did not significantly impact the binding of 2022 (Table 1); in addition, we have also assessed the binding of 2022 to alanine scanning mutants of ten critical residues of RBD, including D405A, K417A, K444A, Y449A, F456A, K458A, F486A, F490A, V503A, Y505A, related to interaction with ACE-2 in ELISA. 2022 bound strongly to all alanine mutants, despite the fact that 2F8, one of the parental antibody of 2022, completely lost binding to variant V503A (data not shown). Furthermore, the F486 mutation has been shown to reduce the effectiveness of casirivimab (F486I, F486V), bamlanivimab (F486V), and etesevimab (F486V) (26), the F490S is one of the two (F490S, L452Q) mutations carried in RBD by lambda variant, yet, either F486A or F490A has no impact on 2022 binding. Together, these results suggest that 2022 could be effective against SARS-CoV-2 for all VOCs currently known, it also suggests that 2022 could potentially be effective for future variants.

**Figure 2:**
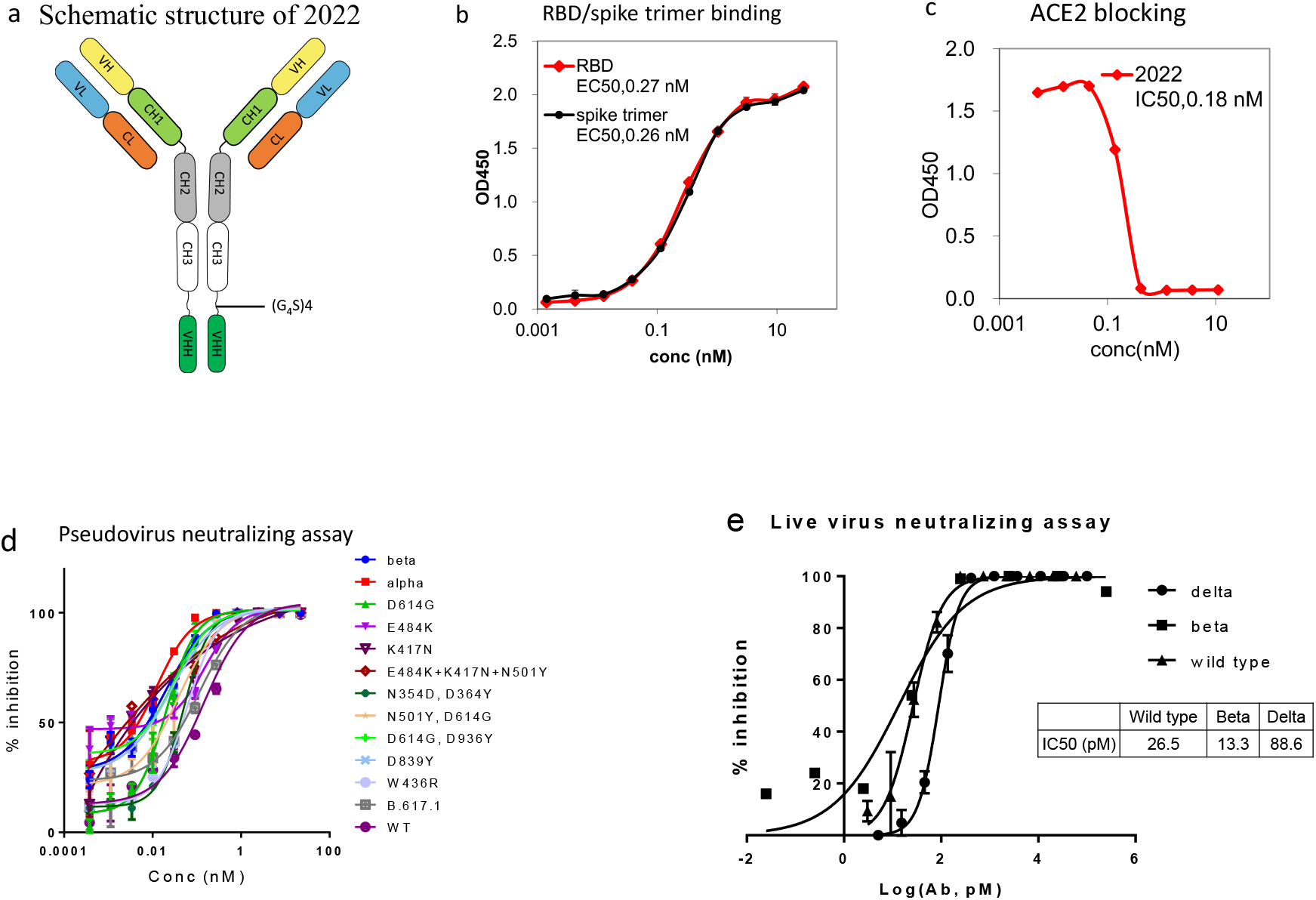
Characterization of 2022 in vitro. a. Schematic structure of 2022. b. Binding of 2022 to recombinant RBD or ectodomain spike trimer in ELISA. Data are mean ± S.D. of technical duplicates. EC50 values are shown on the graph. c, Blocking of RBD binding to recombinant human-ACE2 in ELISA. IC50 values are shown on the graph. d. Neutralization potency on SARS-CoV-2 pseudovirus. Data are mean ± S.D. of technical duplicates. e. Neutralization potency on SARS-CoV-2 live virus. Data are mean ± S.D. of technical duplicates. IC50 values are shown on the graph.

**Table 1.** Binding of 2022 to RBD or S1 variants in ELISA. Natural variants of RBD or S1 protein were tested for 2022 binding by ELISA.

| Mutation | Variant | EC50 (pM) |
| --- | --- | --- |
| wildtype | NA | 270 |
| Spike trimer | NA | 260 |
| HV69-70 deletion, Y144 deletion, N501Y,A570D, D614G,P681H | Alpha | 250 |
| K417N, E484K, N501Y, D614G | Beta, Gamma | 220 |
| L452R, T478K | Delta | 260 |
| L452R, E484Q | B.1.617.1 | 250 |
| W436R | NA | 280 |
| N354D,D364Y | NA | 200 |
| V367F | NA | 260 |
| N501Y | NA | 250 |
| K417N | NA | 250 |
| D614G | NA | 220 |

The binding kinetics of 2022 was measured by surface plasma resonance (SPR). 2022 was immobilized to Protein-A sensor and the affinities to RBD or spike ecto-trimer were measured. Consistent with the results of ELISA, 2022 bound with very high affinity to both the RBD and spike trimer of wild-type SARS-CoV-2, with equilibrium dissociation constant (KD) values of 46 pM and 48 pM, respectively. In addition, 2022 also bound mutant RBD or S1 with mutations found in all major variants of concern reported so far with high affinities, with KD values ranging from 1060 pM to 59 pM, which are 5 to 10-fold better than the benchmark neutralizing antibody 10933 for all variants tested (21) (Table 2), notably, 2022 has an affinity of 18.6 pM for delta RBD.

**Table 2.** 2022 binding kinetics as measured by Surface Plasmon Resonance. KD: equilibrium dissociation constant

|  | antigen | mutation | KD (M) | 19033 |
| --- | --- | --- | --- | --- |
| WT | RBD monomer | NA | 4.59E-11 | 1.03E-09 |
| WT | Spiker-trimer | NA | 4.77E-11 | 4.18E-10 |
| Alpha | S1 monomer | HV69-70 deletion, Y144 deletion, N501Y, A570D, D614G, P681H | 4.02E-10 | 4.70E-09 |
| Beta | S1 monomer | L18F, D80A, D215G, LAL242-244 deletion, R246I, K417N, E484K, N501Y, D614G | 1.06E-09 | 8.66E-09 |
| Gamma | S1 monomer | L18F, T20N, P26S, D138Y, R190S, K417T, E484K, N501Y, D614G, H655Y | 8.10E-10 | 8.20E-09 |
| Delta | RBD monomer | L452R, T478K | 1.86E-11 | 1.27E-10 |
| B.1.617.1 | RBD monomer | L452R, E484Q | 5.97E-11 | 1.86E-09 |

### 2022 potently neutralizes SARS-CoV-2 infection

The neutralization activity of 2022 against SARS-CoV-2 was assessed using both pseudovirus and live virus assays. The neutralization potency of 2022 was tested against luciferase reporter viruses pseudotyped with SARS-CoV-2 spike protein. As shown in Figure 2d and Table 3, 2022 exhibited highly potent neutralization activity against the wild-type and all mutant pseudovirus carrying mutations found in the past and the currently dominant VOC, such as alpha, beta, gamma and delta, with IC50 values ranging from 1 pM to 270 pM, IC90 values from 50 pM to 690 pM, including mutations known to be resistant to etesevimab (k417N) or bamlanivimab (E484K) (26). Notably, 2022 showed exceptional potency for delta pseudovirus, with an IC50 value of 1 pM. The neutralization activity of 2022 was next tested against SARS-CoV-2 live virus by focus reduction neutralization test (FRNT) assay. 2022 showed exceptionally potent neutralization activity against wild-type live SARS-CoV-2, with an IC50 value of 26.4 pM and an IC90 value of 114 pM. 2022 also showed extremely potent neutralizing activity against the beta (IC50 = 13 pM) and delta variants (IC50 = 88.6 pM), (Table 4).

**Table 3.**
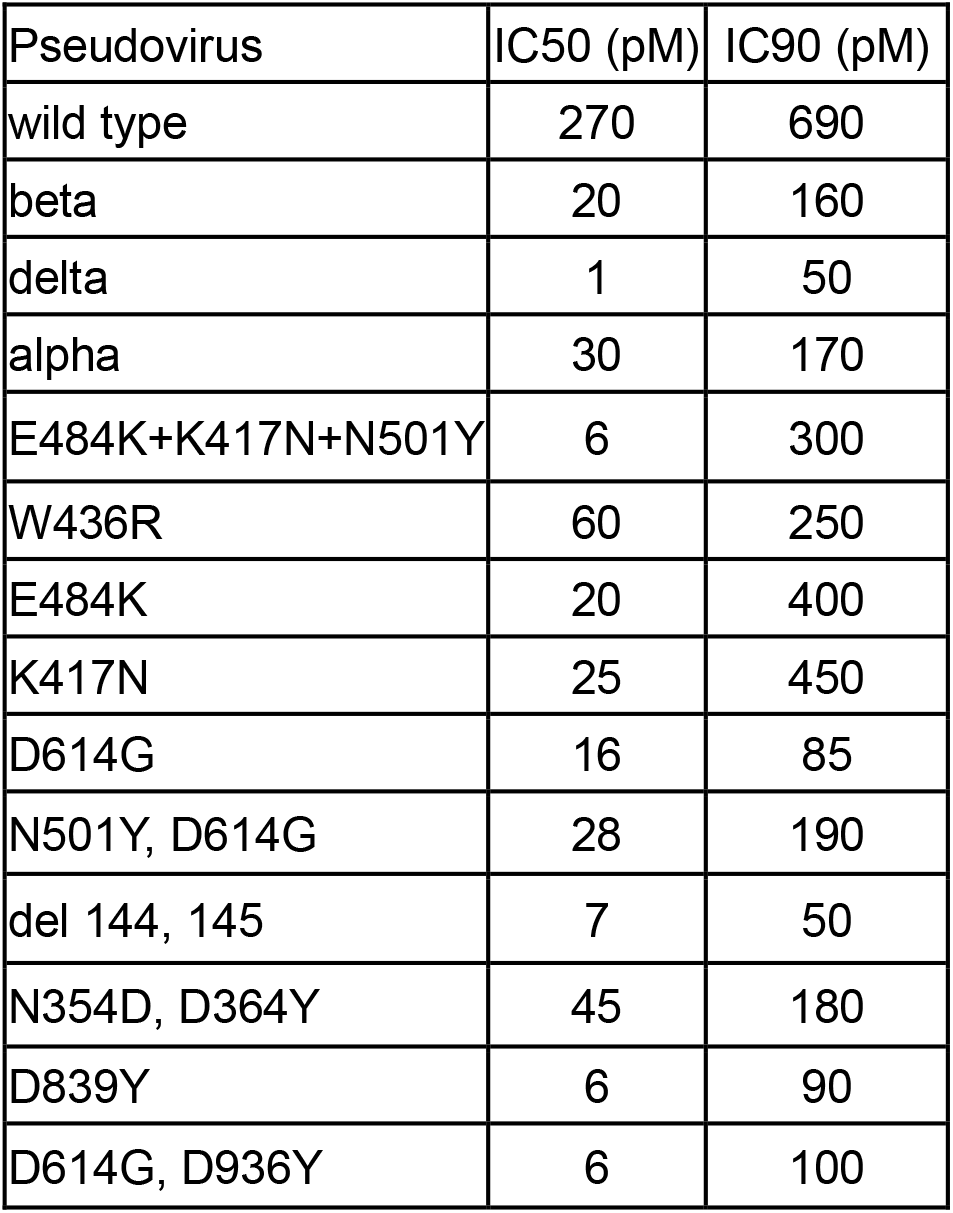
Pseudovirus neutralization of SARS-CoV-2 and its variants. IC50 = concentration inhibiting maximal activity by 50%; IC90 = concentration inhibiting maximal activity by 90%.

| Pseudovirus | IC50 (pM) | IC90 (pM) |
| --- | --- | --- |
| wild type | 270 | 690 |
| beta | 20 | 160 |
| delta | 1 | 50 |
| alpha | 30 | 170 |
| E484K+K417N+N501Y | 6 | 300 |
| W436R | 60 | 250 |
| E484K | 20 | 400 |
| K417N | 25 | 450 |
| D614G | 16 | 85 |
| N501Y, D614G | 28 | 190 |
| del 144, 145 | 7 | 50 |
| N354D, D364Y | 45 | 180 |
| D839Y | 6 | 90 |
| D614G, D936Y | 6 | 100 |

**Table 4.** Neutralization potency of 2022 against SARS-CoV-2 live virus by focus reduction neutralization test (FRNT). Data presented as mean of duplicate wells. IC50 = concentration inhibiting maximal activity by 50%; IC90 = concentration inhibiting maximal activity by 90%.

| Variant | IC50 (pM) | IC90 (pM) |
| --- | --- | --- |
| wild type | 26.4 | 114 |
| Beta | 13.3 | 35.4 |
| Delta | 88.6 | 259 |

### Prophylactic efficacy of 2022 in mouse model of SARS-CoV-2

The in vivo efficacy of 2022 was assessed in a mouse model of SARS-CoV-2. In this model, mice are first transduced intranasally with an adenovirus that expresses human ACE2 (AdV-hACE2), which results in expression of hACE2 in mice alveolar and airway epithelium, leading to the susceptibility of mice to be infected with SARS-CoV-2 live virus (27). To assess the prophylactic efficacy of 2022, animals were treated with a single dose of 1 mg of 2022, or PBS one day before intranasally challenging with 1 × 10^5^ plaque-forming unit (PFU) of SARS-CoV-2 live virus. As previously reported, substantial rapid weight loss of up to 20% at 4 days post infection (dpi) was observed in all 5 animals treated with PBS. By contrast, mice treated with 2022 prophylactically either intranasal (i.n.) or intraperitoneal (i.p) had no significant loss of body weight during the entire experiment (Figure 3b). The virus titers were determined in the whole lung tissue homogenates at 3 dpi by FRNT assay. The virus titers in the lungs were below the detection limit in all mice receiving 2022 prophylactically either i.n or i.p, compared with about 1×10^5^ pfu/g lung in all mice treated with PBS, indicating that 2022 could abolish virus replication in the lungs (Figure 3c). These results indicated that 2022 can block the establishment of virus infection when given prophylactically, suggesting that 2022 is efficacious prophylaxis both i.n and i.p, as shown by undetectable viral load in the lungs and no weight loss.

**Fig. 3.**
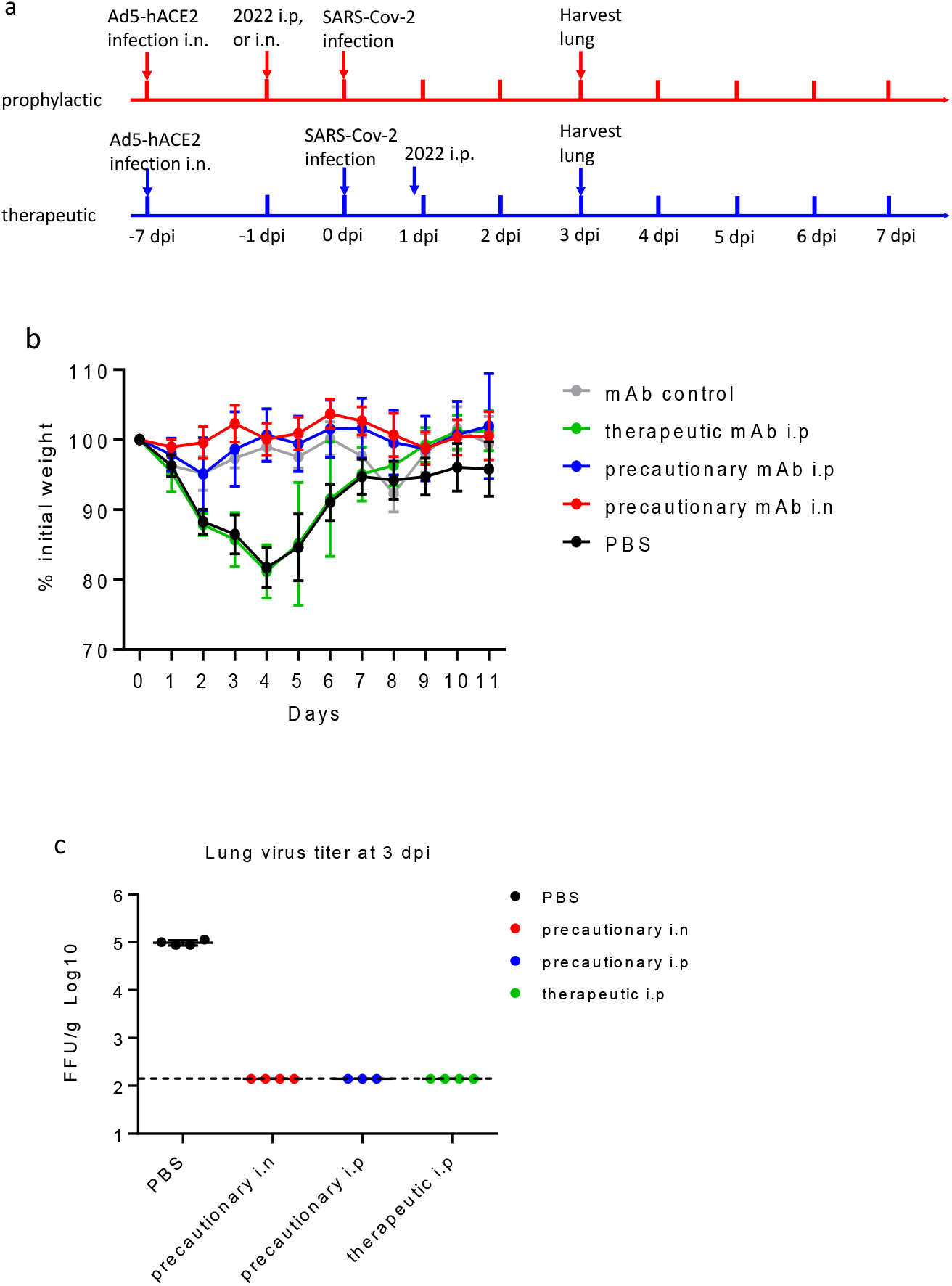
In vivo efficacy of 2022 in mouse model of SARS-CoV-2. a. Schematic overview of study design. Animals were administered i.p. or i.n. a single dose (50 mg/kg) of 2022, or PBS. Animals were treated 24 hours before or 18 hours post virus challenge for precautionary and therapeutic, respectively. Body weight was recorded daily for 11 days. b. Body weight changes of animals. Data are presented as mean ± SD. (n=5). c. Whole lung virus titer at 3 dpi as assessed by FRNT assay. Each dot represents an individual animal. Dotted line indicates the detection limit of the FRNT assay.

### Therapeutic efficacy on mouse model of SARS-CoV-2

The efficacy of 2022 was then tested in the same mouse model in a therapeutic setting. Mice were treated i.p with a single dose (1 mg/mouse) of 2022 18 hours after challenging with 1×10^5^ plaque-forming units (PFU) of SARS-CoV-2 virus. Because viral replication and disease progress occurred very rapidly in mice challenged with 1×10^5^ PFU of virus in this model (27), far exceeding the time course of COVID-19 in clinical, the therapeutic setting in this model represented a very high bar, more closely mimicking late and more severe disease stages in the clinic. Nevertheless, significant therapeutic benefit was observed in mice treated with 2022. The virus titers in the lung at 3 dpi were undetectable in all animals received 2022, as compared with ∼1×10^5^ FFU/lung in all animals treated with PBS (Figure 3c), demonstrating that 2022 can constrain virus replication in vivo even when virus infection was vigorously ongoing. Even though weight loss was not prevented, due possibly to the rapid progress of the disease setting, animals receiving 2022 recovered faster than did PBS-treated mice. 2022-treated mice regained fully their body weight loss by 11 dpi whereas PBS-treated animals did not (Figure 3b). In summary, these in vivo results demonstrate that 2022 is efficacious against SARS-COV-2 infection in mice as prophylaxis or therapeutic treatment, and that 2022 could be promising candidate for the prevention and treatment of COVID-19 induced by SARS-COV-2 wild-type or VOCs,such as delta.

## Discussion

In this study a bi-specific monoclonal antibody, 2022, was derived from two antibodies with distinct non-overlapping epitopes on RBD, which combines the advantages of two antibodies into a single molecule, retains potency and breadth, shows potent RBD-binding and virus neutralizing activity across all of the reported VOCs (alpha, beta, gamma, and delta) and a wide variety of variants with mutations known to compromise the effectiveness of neutralizing antibodies and vaccines currently used in clinic.

2022 has demonstrated potent in vivo efficacy against SARS-CoV-2 in mice both prophylactically and therapeutically, protecting animals from disease progress, abolish virus replication in the lung. When given prophylactically, animals were completely protected from body weight loss. Our results showed that 2022 can prevent infection in an animal model either as i.n. or i.p. It would be interesting to evaluate how long the protection can last. Even though weight loss was not prevented in therapeutic setting but animals receiving 2022 did regain weight loss faster than did placebo-treated animals. Previous study has shown prevention of weight loss in similar mouse model, but it is with significantly less virus load (1×10^4^ PFU vs 1×10^5^ PFU) and earlier dosing time point (12 hours vs 18 hours) post virus challenge (19). These results, together with other’s findings, suggest that disease progress in this model is virus-load and treatment dosing-time post infection sensitive.

Our data suggest that a bi-specific antibody might likely be more effective in blocking host-virus interactions and prevention of viral escape than cocktails of two antibodies. For instance, 2022 showed strong binding to variant even to which its parental antibody did not bind. Physically connecting the two non-over-lapping epitope-binding antibody likely increases overall association constants (Kon values) and decreases overall dissociation constants (Koff values), thus increasing the affinity, and also increasing the avidity of the bi-specific antibody via its increased valency in a single molecule. Previous study has also demonstrated that biparatopic bispecific antibody have significantly higher affinity and stronger neutralizing activity against SARS-CoV-2 (28). Overall, a single molecule with bi-specificity and multi-valency will likely has better efficacy and broader spectrum against mutations. For instance, recently reported monoclonal antibody 2B11 showed potent neutralizing ability against wild-type SARS-CoV-2, yet, its neutralization activity against B.1.351 or P.1 pseudovirus was significantly impaired (29).

In conclusion, 2022 maintains strength and breadth across all of the reported VOCs (alpha, beta, gamma, and delta) and a wide variety of variants with mutations known to compromise the effectiveness of neutralizing antibodies and vaccines currently used in clinic. 2022 would be an attractive solution to the ongoing resurgence of Covid-19 pandemic caused mainly by VOCs, such as delta, and possibly to potential future newly emerging variants and warrants evaluation in human clinical studies.

## Acknowledgement

We are grateful to our colleagues at Bio-Thera Solutions for their support during the study, especially Xiong Mei, Jian Ma, Yanli Liu, Wengrong Huang, Chao Qin, Junjie Huang, Zhencheng Chen, Chuang Li, and Shide Liang. We thank Bert Thomas for reviewing this manuscript.

## Competing interest

Hui Zhang, Haohui Huang, Zhiwei Wang, Jiaping Li, Junyou Chen, Zhiqi Su, Huafei Su, Li Wang, Chunping Deng, shujun Pei, Shenhua Zhu, Chan Li, Haitao Yue, Cuihua Liu, Jinchen Yu, Shengfeng Li, Xianming Huang are employee and/or stockholders of Bio-Thera Solutions.

## Materials and methods

### Cell lines

HEK293 (ACS-4500TM, ATCC) and African Green monkey kidney-derived VeroE6 cells (CRL-1587, ATCC) were cultured and passaged in DMEM with 10% FBS. HEK293-hACE2 was constructed and sorted by Bio-Thera Solutions, Ltd.

### Recombinant Proteins

Biotinylated 2019-nCoV S protein RBD, His,Avitag™ (SPD-C82E9,Acrobiosystems); SARS-Vov-2 S protein RBD, His Tag (SPD-S52H6, Acrobiosystems); SARS-CoV-2 S1 protein, His Tag (S1N-S52H5, Acrobiosystems); SARS-CoV-2 S protein (R683A, R685A), His Tag, active trimer (SPN-C52H8, Acrobiosystems); SARS-CoV-2S protein RBD (N354D, D364Y), His Tag (SPD-S52H3,Acrobiosystems); SARS-CoV-2S protein RBD (W436R), His Tag (SPD-S52H7, Acrobiosystems); SARS-CoV-2 S protein RBD (V367F), His Tag(SPD-S52H4, Acrobiosystems); SARS-CoV-2 Spike RBD (L452R, E484Q) Protein (His Tag)(40592-V08H8,Sino Biological);

### Pseudovirus strains

Wildtype pseudovirus (GM-0220PV07, Genomeditech); Variant (D614G) (GM-0220PV14, Genomeditech);Variant (E484K)(GM-0220PV35, Genomeditech); Variant (D614G, D936Y) (GM-0220PV19, Genomeditech); Variant (D839Y) (GM-0220PV6, Genomeditech); Variant (V483A) (GM-0220PV17, Genomeditech); Variant (D614G, A831V) (GM-0220PV24, Genomeditech); Variant (W436R) (GM-0220PV26, Genomeditech); Variant (B.1.1.7/VUI-202012/01 del 145Y) (GM-0220PV33, Genomeditech); Variant (B.1.351/501Y.V2) (GM-0220PV32, Genomeditech).

### Mice

Specific pathogen-free 6-10 weeks old female BALB/c mice were purchased from Hunan SJA Laboratory Animal Co. (Hunan, China) and bred in Animal Care Facilities of Guangzhou Customs Technology Center and Guangzhou Medical University.

### SARS-CoV-2 viruses

The Accession number of wild-type strain used in the research is a clinical isolated SARS-CoV-2 virus 2019-nCoV/IQTC01/human/2020/Guangzhou (GenBank : MT123290.1). The SARS-CoV-2 beta variant (SARS-CoV-2/human/CHN/20SF18530/2020 strain was isolated from an infected male individual and passaged on Vero E6 cells; Accession number on National Pathogen Resource Center is NPRC2.062100001) The SARS-CoV-2 delta variant was a gift from Guangdong Provincial Center for Disease Control and Prevention.

### Panning of phage display library

Our proprietary fully-synthetic Intelligent Designed and Engineered Antibody Libraries (IDEAL) contains one VH/VL-display phage library and one VHH-display phage library. The two libraries were panned separately on biotinylated recombinant SARS-CoV-2 RBD (SPD-C82E9, Acrobiosystems), for 4 rounds with decreasing amount of antigen, 10 ug, 2.5 ug, 0.5 ug and 0.1 ug of RBD for round 1 to 4, respectively. The phage-RBD complexes were captured with Dynabeads MyOne Streptavidin T1. Prior to each round of panning, the Dynabeads and phage libraries were blocked by 3-5% BSA in PBS. The libraries were incubated with RBD overnight at 4°C, slowly rotating. Then Dynabeads were added into the phage-RBD mixtures, subsequently washed with PBS containing 0.05% Tween 20 and twice with PBS. The remaining phages were eluted with 1 mg/ml trypsin (Sigma-Aldrich) for 30 minutes at room temperature, accompanied by gently rotating. The eluted phages were infected into exponentially growing TG1 Escherichia coli for 30 minutes at 37°C, and amplified to 40 ml 2xYT containing 100 ug/ml ampicillin. The temperature was switched from 37°C to 28°C, overnight shaking at 230 rpm to obtain the amplified phages of each round. After 4 rounds of panning, VH/VL and VHH DNA sequences were pool transferred from the phage displaying vector to the prokaryotic expression vector, then transfected into BL21 Escherichia coli with electroporation. ELISA was carried out on soluble scFv or VHH for RBD-binding.

### Enzyme-linked immunosorbent assay (ELISA)

One ug/ml His-tagged Spike RBD protein of SARS-CoV-2 as well as mutated S1 domains, RBDs, or ectodomain of trimeric Spike were immobilized onto 96-well plates (9018, Corning) overnight at 4°C, plates were blocked with 3% BSA in PBST for 2 hours at 37°C. Samples were serially three-fold diluted and added 100 ul per well into blocked plates, incubated for 1 hour at 37°C. Bound antibodies were detected with Peroxidase conjugated goat anti-human kappa light chains antibody (A7164-1ML, Sigma). 100 ul TMB(Tetramethylbenzidine,Biopanda TMB-S-001) were added per well to develop. Experiments were conducted in duplicates, value=Mean±SD.

### Receptor-binding blocking ELISA

One ug/ml His-tagged spike RBD protein of SARS-CoV-2 (SPD-C52H3, Acrobiosystems) were immobilized onto 96-well plates (9018, Corning) overnight at 4°C, plates were blocked with 3-5% BSA in PBST for 2 hours at 37°C. Serially threefold diluted 2022 was added into the blocked plate, incubated for 1 hour at 37°C. Biotinylated human ACE2 protein (AC2-H82E6, Acrobiosystems) was added to the plate with antibody diluted inocula to the final concentration of 25 ng/ml, further incubated for 1 hour at 37°C. The remaining biotinylated human ACE2 binding to the RBD coating on the plate was detected using HRP-labelled strepavidin (016-030-084, Jackson Immunoresearch). The plate were developed with TMB (Tetramethylbenzidine,Biopanda TMB-S-001). Absorbance at 450 nm was measured on SpectraMax Plus Absorbance Microplate Reader (Molecular Device, CA).

### Antibody and recombinant RBD expression and purification

The 2022 coding sequences of heavy and light chains were cloned to expression plasmids, respectively. HEK293 cells were transiently co-transfected at a ratio of 1:2 (H:L) with PEI (49553-93-7, Polyscience) according to the manufacturer’s instruction. The supernatants were harvested at day 7 post-transfection and purified by protein-A affinity column. For expression of recombinant alanine mutants, we designed and synthesized 10 RBD DNA sequences (GENERAL BIOL), each one contained one specific alanine mutant of 10 critical residues related to interaction with ACE2, and cloned into expression vector with Fc tag. Transiently transfected to HEK293 cells to obtain recombinant RBD alanine mutants and purified by protein A columns.

### Affinity measurement of antibodies by Surface plasmon resonance (SPR)

SPR experiments were all conducted with a Biacore T200 system (GE Healthcare); All assays were performed with a Sensor Chip Protein A (GE healthcare),with a HBS EP+ running buffer (0.1M HEPES, 1.5M NaCl, 0.03M EDTA supplemented with 0.005% vol/vol surfactant P20 at 25°C.) To determine the affinities of nanobody VHH18, human IgG1 antibody 2F8, and 2022 to SARS-CoV-2 RBD-His tag, spike trimer-His tag and other S1-His tag variants, antibodies were immobilized onto the sample flow paths of the sensor Protein A chip. The reference flow cell was left blank. SARS-CoV-2 RBD-His tag, or spike trimer-His tag or other S1-His tag variants was injected over the above-mentioned flow paths at a range of five concentrations prepared by two-fold serial dilutions started at 50nM, at a flow rate of 30 ul/min, with an association time of 75s and a dissociation time of 180s. HBS EP+ running buffer was also injected using the same program for background subtraction. All the data were fitted to a 1:1 binding model using Biacore T200 Evaluation Software 3.1.

### Epitope binning by in-tandem fortebio assay

Epitope analysis for 2F8 and VHH18 was carried out by in-tandem biolayer interferometry (BLI) via Octet QKe (ForteBio) according to the manufacturer’s instruction. His, Avitag SARS-CoV-2 RBD protein (SPD-C82E9, Acrobiosystems) was diluted to a final concentration of 400 nM in kinectics buffer (1×PBS, 0.01% Tweens-20) and immobilized onto Strepavidin biosensor. The sensor was saturated with the first antibody, either 2F8 Fab or VHH18-His, subsequently, the above bioprobes were flown over with the different second antibody, either VHH-his or 2F8 Fab, respectively.

### Pseudovirus neutralization assay

HEK293 was transfected with human ACE2 cDNA cloning vector (HG10108-M, Sino Biological), and sorted with BD FACJazz cell sorter to get monoclonal cell line, HEK293-hACE2. 2022 was serially threefold diluted and incubated with SARS-CoV-2 pseudovirus or other variant pseudoviruses which were diluted according to manufacturer’s instruction for 1 hour at 37°C. HEK293-hACE2 was detached and then added to the pseudovirus-antibody mixtures. After 48 hours incubation at 37°C in 5% CO_2_, neutralization potencies were quantified in luciferase assay measured with Bio-Lite Luciferase Assay solutions (DD1201-03, Vazyme). The values were read on microplate reader SpectraMax M3 (Molecular Devices). The half maximal inhibitory concentration (IC50) and 90% of maximal inhibitory concentration (IC90) were determined by four-parameter logistic regression. Experiments were performed in duplicate.

### Focus reduction neutralization test (FRNT)

2022 was serially threefold diluted in DMEM, and incubate with an equal volume of SARS-CoV-2 wildtype or delta variant containing 200 PFU for 1 hour, at 37°C. The mixtures were then added to Vero E6 monolayers in a 96-well plate in triplicate and incubate for 1 hour at 37°C in 5% CO2. The inocula were removed, and added 100ul pre-warmed 1.6% (w/v) CMC (carboxylmethylcellulose) in MEM containing 2% FBS per well, further incubate for 24 hours at 37°C in 5% CO2. Cells were then fixed with 4% paraformaldehyde (PFA) and permeabilized using 0.2% Triton X-100. Cells were tested with a rabbit anti-SARS-CoV-2 nucleocapsid protein polyclonal antibody (Cat. No.: 40143-T62, SinoBiological, Inc.), and an HRP-labelled goat anti-rabbit as secondary antibody (111-035-003, Jackson ImmunoResearch). The foci were developed by Trueblue peroxidase substrate and results were read on Immuno-ELISPOT (CTL ImmunoSpot UV). Values were determined using four-parameter logistic regression (GraphPad Prism). Experiments were conducted under the standard operating procedures of the approved Biosafety Level-3 laboratory.

### Animal experiment

All animal experiments were performed under the relevant ethical regulations regarding animal research. The mice experiments for in vivo efficacy were conducted in the Animal Care Facilities of Guangzhou Customs District Technology Center and Guangzhou Medical University. All protocols were approved by the Institutional Animal Care and Use Committees of Guangzhou Customs District Technology Center and Guangzhou Medical University. All the live virus experiments were conducted in the Biosafety Level-3 laboratory of Guangzhou Customs District Technology center following standard operating procedures.

Balb/c mice were mildly anesthetized with isoflurane and i.n. transduced with 2.5×10^9^ PFU of Ad5-ACE2 in 75ul DMEM. Four days after Ad5-ACE2 transduction, mice were infected i.n. with 1×10^5^ PFU SARS-CoV-2 virus. For prophylaxis, mice were dosed with 1 mg antibody either i.p or i.n. 24 hours before virus infection; for therapeutic treatment, mice were dosed with 1 mg antibody i.p. 18 hours post virus challenge, PBS was used as placebo. Animals were monitored daily and body weight were recorded daily. Lungs tissue were harvested at 3 dpi for virus titers measurement by focus reduction neutralization test (FRNT). About 140 mg of lung tissue from each mouse (n=3, or 4) was homogenized to 1 ml PBS, 50 ul lung homogenate supernatants were used for virus titers.

